# Genetic diversity, haplotype analysis, and risk factor assessment of hepatitis a virus (HAV) isolates from the West Bank, Palestine during the period between 2014 and 2016

**DOI:** 10.1101/2020.09.25.313015

**Authors:** Kamal Dumaidi, Hayah Qaraqe, Amer Al-Jawabreh, Rasmi Abu-Helu, Fekri Samarah, Hanan Al-Jawabreh

## Abstract

**Background:** HAV genotypes and its genetic diversity is rarely investigated in our region as well as worldwide.

**Aims:** the aims of the present study were to determine the HAV genotypes and its risk factors and to investigate the genetic diversity of the HAV isolates in the West bank, Palestine.

**Study design:** a cohort of 161 clinically and laboratory confirmed HAV (IgM-positive) cases and 170 IgM negative individuals from all the districts of the West Bank, Palestine during the period of 2014-2016 were tested for VP3/VP1 junction of the HAV genome using RT-PCR and sequence analysis. Phylogenetic analysis, genetic diversity and haplotypes analysis were used to characterize the VP3/VP1 sequences.

**Results:** Overall, all the 34 sequences of the HAV was found to be HAV-IB sub-genotype. The phylogenetic analysis showed four main clusters with cluster III exclusively consisting of 18 Palestinian isolates (18/23-78%) with weak bootstrap values. A high haplotype diversity (Hd) and low nucleotide diversity (π) were observed. Cluster III showed high number of haplotypes (h=8), but low haplotype (gene) diversity (Hd=0.69). A total of 28 active haplotypes with some consisting of more than one sequence were observed using haplotype network analysis. The Palestinian haplotypes are characterized by closely related viral haplotypes with one SNV away from each other which ran parallel to cluster III in the phylogenetic tree. A smaller Palestinian haplotype (4 isolates) was three SNVs away from the major haplotype cluster (n=10) and closer to haplotypes from Iran, Spain, and South Africa. Young age, low level of parent’s education, poor hand washing and drinking of un-treated water was considered the major HAV risk factors in the present study.

**Conclusion:** HAV-IB subgentype is endemic in Palestine. HAV showed low genetic variation and nucleotide diversity. Furthermore, haplotype network analysis revealed haplotype variation among the Palestinian sequences.

## Introduction

Hepatitis A virus (HAV) is “a nonenveloped RNA virus belonging to the family Picornaviridae, genus Hepatovirus”. HAV is one of the major causes of acute hepatitis worldwide and contributes to substantial morbidity in both developed and developing countries. Based on HAV genome sequences, human HAV have been classified into three genotypes, HAV I, II, III and sub-divided into 6 sub-genotypes (IA, IB, IIA, IIB, IIIA, IIIB) [1].

Worldwide, the incidence rate of the HAV infection is underestimated due to the clinical presentations of this disease, since infection at early childhood is largely passed asymptomatic and/or has mild forms [2]. Recent data showed that the global incidence of HAV is 1.9% with an estimate of 119 million cases of HAV infection [3, 4]. Approximately, 1.4 million new cases of HAV reported each year with up to 22% of the cases being hospitalized [5]. Palestine (West Bank, and Gaza Strip) was classified as an area of very highly endemic area [6]. The Palestinian official figures put the HAV infection incidence rate as high as 9.5-85 per 100,000 during the period between 2000 and 2018 [7]. Actual incidence rate is thought to be higher due to underreporting and asymptomatic cases.

HAV is transmitted mainly by the fecal-oral route and mainly through drinking and consumption of contaminated water and food and to a lesser extend from person to person or via blood transfusion [8–10]. Poor hygiene and sanitation practice reported as the major risk factors for HAV infection, particularly in low and middle-income countries [11].

The World Health Organization (WHO) classified the HAV endemicity based on anti-HAV IgG antibodies as follows: high (≥ 90% IgG seroprevalence by 10 years of age), intermediate (≥ 50% IgG seroprevalence by 15 years of age, < 90% IgG seroprevalence by 10 years of age), and low (≥ 50% IgG seroprevalence by 30 years of age, < 50% IgG seroprevalence by 15 years of age) [12].

The viral infection is characterized by asymptomatic or mild in children at early age of life. On the contrary, adult infections are more frequently occurring with symptoms. In high endemic regions, HAV infection is acquired in early age of childhood and most adult is positive for anti-HAV IgG with a life-long immunity, whereas, in low endemic countries, most adult population is susceptible to infection[3]

A recent review investigated and analyzed data based on anti-HAV seroprevalence in the Middle East and North Africa (MENA) countries reported a gradual shift in the age of HAV infection from early childhood to late child and adult-hood and indicating a shift towards intermediate endemicity in these countries in general [6]. However, based on only one orphan, old study conduct in Gaza between 1995 and 2001, Palestine is still considered a high endemic country. Therefore, no solid conclusion has been made regarding the current endemicity levels of HAV and vaccine recommendations in Palestine [13]

The aims of this study were to determine the HAV genotypes, the risk factors associated with HAV infections and to visualize the genealogical relationship between intraspecific HAV individual genotypes at the population level and exploring the genetic diversity between the HAV sequences.

## Materials and methods

### Study sample

This case-control, cross-sectional study design, comprised of 161 clinically and laboratory confirmed HAV (IgM-positive) cases and 170 apparently healthy controls from all the districts of the West Bank, Palestine during the period of 2014-2016. A special questionnaire was used to collect demographic and medical data including age, sex, residence, education, working status, housing details, toilet facilities (flush vs. pit), drinking water sources (pipe-public net sources vs. collected well or local spring), available used sewage systems (public net system vs. holes), income, educational level of parents, and other socio-economic data. The study was approved by the Palestinian Ministry of Health under the reference number 145/1541/2014. Verbal informed consent was obtained from all patients or their guardians in case of minors. The data was analyzed anonymously.

### Serological assays

All serum samples of both the HAV cases and the healthy subjects were tested for HAV IgM antibodies at the Central Public Health Laboratories, Palestinian Ministry of Health using commercially IgM capture ELISA (Architect, Abbot-USA, ELISA) with a sensitivity and specificity of >99% according to the manufacturer’s instructions.

### Molecular assays

#### Extraction of viral RNA

The HAV viral genome was extracted from 200μl of serum samples, using a QIAamp Mini Elute Virus spin kit for the viral RNA/DNA extraction (QIAGEN, Germany) according to the manufacturer’s instructions, and kept at −20C until testing.

#### Reverse transcriptase-polymerase chain reaction (RT-PCR)

The HAV genomes of 136 serum samples from IgM-positive patients were amplified using two primers targeting the VP3/VP1 region of the viral genome as described previously by Lee *et al.*, (2012) [14]. Briefly, the synthesis of the cDNA and the RT-PCR was carried out in 25μl reaction mixture containing 4μl viral RNA extraction, 10U Reverse transcriptase (AMV), 10pmol of each the forward and the reverse primers: (HAV1; 5’ - GCTCCTCTTTATCATGCTATGGAT-3’ and rHAV2; 5’-CAGGAAATGTCTCAGGTACTTTC-3’) and 12.5μl of PCR Reddy master mix (Thermo Scientific). PCR products (6μl) were loaded onto a 2% agarose gel, electrophoresed, and stained with ethidium bromide for band visualization at an expected length of 244bp using the Gel Doc System 2000 (Bio-Rad Laboratories-Segarate, Milan, Italy). Of the 136 PCR-positive sample, 34 representative PCR amplicons were selected randomly for sequence analysis. The PCR amplicons of the 34 samples were purified and sequenced. The HAV identity search was conducted using GenBank Basic Local Alignment Search Tool (BLAST) http://www.ncbi.nlm.nih.gov/BLAST/Blast.cgi.

#### Genetic diversity analysis

The 23 sequences generated by the study and the 28 sequences retrieved from the Genbank were aligned using the MEGA version X [15]. Maximum likelihood phylogenetic tree with 1000 iterations for bootstrapping was constructed using MEGA version X.

Population nucleotide diversity indices such as nucleotide diversity per site (π), average number of nucleotide differences (k), mean genetic diversity (Hd), genetic differentiation parameters (Fst and Nm) and neutrality tests including Tajima’s D and Fu Li’s F test were calculated using DnaSP ver. 6.12.03 [16]. The PopArt 1.7 [17] was used to construct a median-joining haplotype network analysis based on country of source of viral genomes to estimate relationship between haplotypes using nexus input files produced by DnaSP version 6.12.03. Haplotype network analysis was colored and edited using a free open-source vector graphics editor called Inkscape 1.0 (www.inkscape.org). The haplotype analysis was double checked by reconstructing the median joining tree using Network 10 (https://www.fluxusengineering.com/) with RDP input file generated by DnaSP ver. 6.12.03 and default parameters of the software including epsilon value of zero and the connection cost method of Röhl for genetic distance calculation [17]. Greedy FHP method for the genetic distance calculation was also used [18].

### Risk assessment analysis

The EpiInfo, a free CDC statistical package, was used to analyze data. Odds ratio, 95% confidence interval, and Fisher’s exact test were used to assess the risk factors of HAV infection. P-value was considered significant when less than 0.05.

## Results

### Characteristics of study population

A total of 331 individuals were included in the study with a median age of 15 years and a 1:1 female-to-male ratio. The HAV case group were from all 11 Palestinian districts in the West Bank-Palestine with 35% (56/161) from the Al-Khalil district. All the 34 sequenced samples were shown to be of HAV-IB genotype. Twenty-three sequences were deposited in the gene Bank (Table 1)

**Table 1.** Demographic data of the sequenced HAV patient samples

| Accession No. | Sample<br>code/year of<br>isolation | genotype | Age (yr) | District |
| --- | --- | --- | --- | --- |
| MN744241 | 1B/Pal-2014 | HAV-IB | 28 | Bethlehem |
| MN744242 | 11B/Pal-2014 | HAV-IB | 15 | Bethlehem |
| MN744243 | 104S/Pal-2016 | HAV-IB | 6 | Salfit |
| MN744244 | 41K/Pal-2015 | HAV-IB | 14 | Al-Khalil |
| MN744245 | 44Q/Pal-2015 | HAV-IB | 14 | Qalqilia |
| MN744246 | 46N/Pal-2015 | HAV-IB | 25 | Nablus |
| MN744247 | 73B/Pal-2015 | HAV-IB | 2 | Bethlehem |
| MN744248 | 75B/Pal-2015 | HAV-IB | 16 | Bethlehem |
| MN744249 | 86K/Pal-2016 | HAV-IB | 7 | Al-Khalil |
| MN744250 | 98T/Pal-2016 | HAV-IB | 9 | Tulkarem |
| MN744251 | 6B/Pal-2016 | HAV-IB | 8 | Bethlehem |
| MN744252 | 8K/Pal-2014 | HAV-IB | 6 | Al-Khalil |
| MN744253 | 16K/Pal-2015 | HAV-IB | 4 | Al-Khalil |
| MN744254 | 25K/Pal-2016 | HAV-IB | 7 | Al-Khalil |
| MN744255 | 26K/Pal-2016 | HAV-IB | 8 | Al-Khalil |
| MN744256 | 32K/Pal-2016 | HAV-IB | 8 | Al-Khalil |
| MN744257 | 33K/Pal-2016 | HAV-IB | 13 | Al-Khalil |
| MN744258 | 44Q/Pal-2016 | HAV-IB | 14 | Qalqilia |
| MN744259 | 60R/Pal-2016 | HAV-IB | 19 | Ramallah |
| MN744260 | 66K/Pal-2016 | HAV-IB | 8 | Al-Khalil |
| MN744261 | 88K/Pal-2016 | HAV-IB | 12 | Al-Khalil |
| MN744262 | 89N/Pal-2016 | HAV-IB | 5 | Al-Khalil |
| MN744263 | 69T/Pal-2015 | HAV-IB | 21 | Tulkarem |

### Phylogenetic analysis

Phylogenetic tree of the VP3/ VP1 junction region of HAV genome was conducted using the 23 HAV sequences from strains isolated from Palestinian patients along with 28 sequences retrieved from the Genbank. The phylogenetic tree showed four main clusters with cluster III exclusively consisting of 18 Palestinian isolates. Cluster I consisted of 12 strains, of which four were isolated in Palestine (Figure 1). However, the four clusters were weakly supported by bootstrap values (Fig 1).

**Figure 1.**
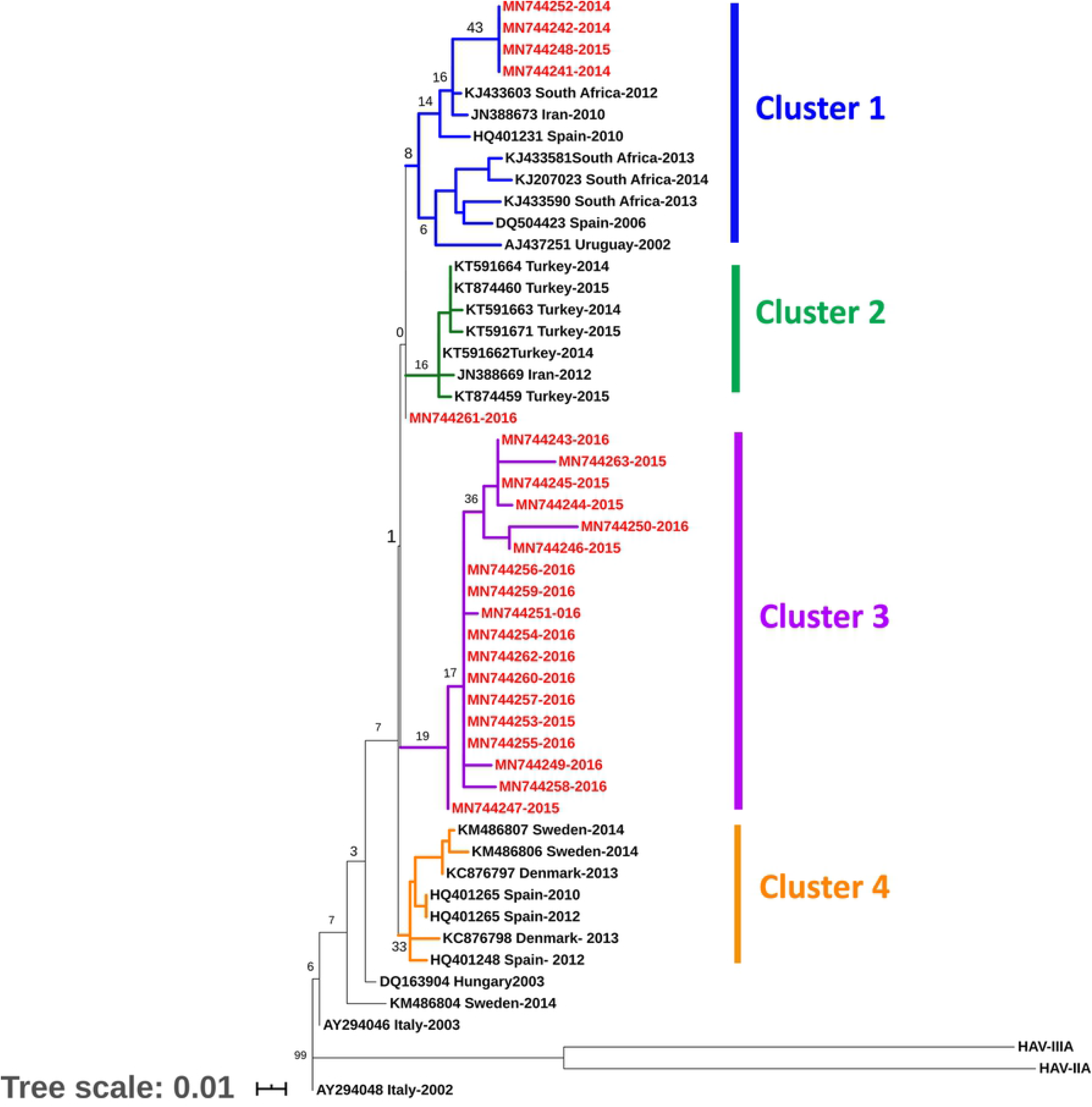
Consensus Maximum Likelihood phylogenetic tree (1000 replicates) based on VP3/VP1 region. The percentage of trees in which the associated taxa clustered together is shown next to the branches. The branch labels in red represent the study sequences, while the black labels represent the sequences retrieved from the gene bank. The different color branches depict different clades. The tree is drawn to scale, with branch lengths measured in the number of substitutions per site. This analysis involved 51 nucleotide sequences conducted in n MEGA X [15, 19].

### Genetic differentiation and Diversity

Population nucleotide diversity indices and neutrality tests were calculated for the VP3/VP1 junction region of the HAV genome, based on phylogenetic clusters (Tables 2 and 3). A high haplotype diversity (Hd) and low nucleotide diversity (π) were observed (Table 2). The total haplotype (gene) diversity (Hd) for the 23 HAV sequences from Palestine and 28 from the Genbank was 0.93± 0.07. At the same time, the total nucleotide diversity per site (π) was 0.01± 0.002, confirming low genetic diversity in the HAV study bulk. The average number of nucleotide differences between any two sequences (k) was 1.7 which is very low. The DnaSP ver. 6.12.03 estimated the total number of haplotypes for the four probable cluster at 24 with highest in cluster I (h=9) (Table 2). Cluster III is composed of 18 isolates, exclusively Palestinian isolates, while one third of cluster I was from Palestine. The Palestinian cluster (III) showed high number of haplotypes (h=8), but lowest haplotype-to-sequence (h:n) ratio (0.4:1), compared to the other clusters and lowest haplotype (gene) diversity (Hd=0.69). Haplotype diversity (H_d_), and number of segregating (polymorphic) sites (S) were highest in cluster I which included Palestinian strains (4/12) along with those from other countries (Figure 1); confirming the highest level of genetic diversity between all probable clusters. Nucleotide diversity (π) is equally low in all clusters (0.01± 0.002). The average number of nucleotide differences (k) is also equal in all clusters except cluster I. Tajima’s D and Fu-Li’s F tests were negative for cluster I, II, and III. The negative values of Tajima’s D and Fu-Li’s, though not statistically different from neutral expectations supported the low population differentiation. Cluster I and III had relatively the highest negative values for Tajima’s D and Fu’Li’s F statistics. However, it did not depart significantly from neutrality (P>0.01) (Table 2). Cluster IV recorded a positive Tajima’s D value (0.17), but showing no statistically significant departure from neutrality (P>0.01) (Table 2).

**Table 2.** Haplotype/nucleotide diversity and neutrality tests of the four probable clusters of HAV as calculated for the VP3/VP1 gene

| Cluster | Haplotype- nucleotide diversity |  |  |  |  |  |  | Neutrality tests |  |
| --- | --- | --- | --- | --- | --- | --- | --- | --- | --- |
| | n | h | h:n | $H_d \pm SD$ | $\pi \pm SD$ | K | S | Tajima's $D$ | Fu-Li's $F$ |
| Cluster-I | 12 | 9 | 0.8:1 | $0.90 \pm 0.08$ | $0.01 \pm 0.002$ | 3.3 | 14 | -1.19 | -1.59 |
| Cluster -II | 7 | 4 | 0.6:1 | $0.71 \pm 0.18$ | $0.01 \pm 0.002$ | 1.1 | 3 | -0.30 | -0.51 |
| Cluster-III | 18 | 8 | 0.4:1 | $0.69 \pm 0.13$ | $0.01 \pm 0.001$ | 1.1 | 7 | -1.41 | -1.72 |
| Cluster-IV | 7 | 4 | 0.6:1 | $0.81 \pm 0.10$ | $0.01 \pm 0.002$ | 1.1 | 3 | 0.17 | -0.03 |
| **Total** | **44** | **24** | **0.5: 1** | **$0.93 \pm 0.07$** | **$0.03 \pm 0.002$** | **3.2** | **23** | **-2.63\*** | **-5.16\*\*** |

n: Number of sequences, h: Number of Haplotypes, Hd: Haplotype (gene) diversity, $\pi$ : Nucleotide diversity (per site)[20], K: Average number of nucleotide differences between two randomly chosen sequences from within in the population[21], S: Number of variable/segregating sites.(1 outgroup, 4 have no clear cluster). \*: $P<0.01$ . \*\*: $P<0.02$

**Table 3:** Gene flow and genetic differentiation indices between the four HAV probable clusters estimated from VP3/VP1gene sequences

| Pop 1 | Pop 2 | Fst | Nm | Kxy | Dxy | Gst | Da |
| --- | --- | --- | --- | --- | --- | --- | --- |
| Cluster-I | Cluster-II | 0.46 | 0.12 | 4.10 | 0.021 | 0.098 | 0.009 |
| Cluster -I | Cluster –III | 0.52 | 0.12 | 4.75 | 0.024 | 0.108 | 0.013 |
| Cluster-I | Cluster-IV | 0.52 | 0.12 | 4.61 | 0.023 | 0.073 | 0.011 |
| Cluster-II | Cluster-III | 0.52 | 0.12 | 2.41 | 0.012 | 0.147 | 0.007 |
| Cluster-II | Cluster-IV | 0.53 | 0.12 | 2.45 | 0.012 | 0.135 | 0.007 |
| Cluster-III | Cluster-IV | 0.63 | 0.11 | 3.19 | 0.016 | 0.136 | 0.011 |

Fst: Wright's F-statistics, pairwise genetic distance[22], Nm: Gene flow and population migration among populations [22, 23], Kxy: Average proportion of nucleotide differences between populations. Dxy: The average number of nucleotide substitutions per site between populations[20], $D_a$ : The number of net nucleotide substitutions per site between populations [20], $G_{st}$ : Genetic differentiation index based on the frequency of haplotypes [24].

Inter-population pairwise genetic distance (Fst) between the four HAV probable populations ranged from 0.46 to 0.65 with Nm value from 0.11 to 0.12 (Table 2) indicating genetic differentiation and minimal migration and gene flow (Nm) between subpopulations. Fst for cluster III which is purely Palestinian compared to clusters I, II and IV were highest (0.52, 0.52 and 0.63, respectively). However, genetic differentiation among subpopulation is generally low especially between clusters I and IV (Gst=0.073). The genetic differentiation between clusters is low as supported by other low genetic differentiation parameters including Gst, Da, and Dxy (Table 3).

### Haplotype network analysis

The median-joining haplotype network constructed by PopArt 1.7 using the 49 taxa produced a total of 28 active haplotypes with some of which consisting of more than one sequence (2-10 sequences) (Figure 2a). Peripheral haplotypes mainly had single nucleotide variation (SNV) from central haplotypes (Figure 4a). The Palestinian haplotypes (red circles) are characterized by closely related viral haplotypes with one SNV away from each other. The Palestinian haplotypes (red) formed a cluster of nodes (n=9) surrounding a major node (haplotype) consisting of 10 identical HAV sequences which ran parallel to cluster III in the phylogenetic tree (Figure 1). A smaller Palestinian HAV haplotype consisting of four sequences was three SNVs away from the major haplotype cluster (n=10) and closer to haplotypes from Iran, Spain, and South Africa which again matches cluster I in the phylogenetic tree.

**Figure 2.**
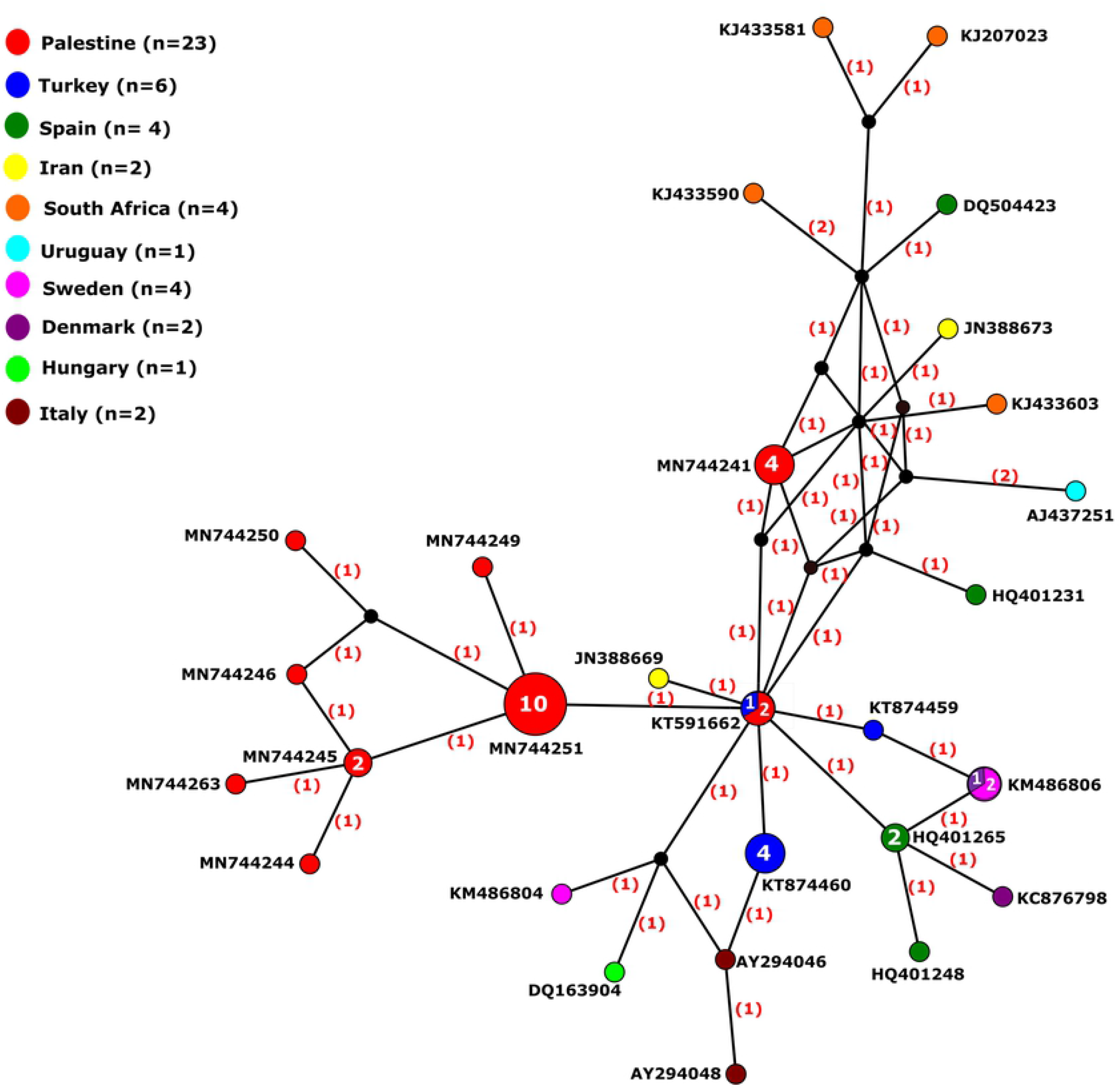
(a) Median-joining haplotype network of HAV viral haplotypes constructed using PopArt 1.7. (b) Median-joining haplotype network reconstructed using Network 10. The network analysis included 28 multiple viral haplotypes of HAV from 2014-2018. Each circle represents a unique haplotype, color represents country of genome origin, and the size of the circle is proportional to number of viral genomes included. The numbers in bracket in (a) represent the number of single nucleotide variations (SNVs) between haplotypes. The lines on the branches in (b) represent the mutated position with one line per mutation

The reconstruction of median joining haplotype network analysis using Network 10 revealed similar haplotype network analysis profile by both distance calculation methods, the Röhl method and the Greedy HPF method (Figure 2b).

### Risk Assessment

The overall, 331 participants (161 cases and 170 health individuals) were analyzed for the HAV risk factors (Table 6). Young age as a demographic variable was found to be significantly associated with HAV infection (OR=7.16, CI: 4.41-11.63, P<0.0001). Level of parents’ education was the only socioeconomic risk factor significantly associated with HAV infections (OR=4.72, CI: 2.64-8.41, P < 0.0001). Furthermore, treating drinking water and washing hands before meal as hygiene and behavior risk variables were found to be statistically significant (R=0.11, CI: 0.01-0.965, P< 0.03 and R= 0.24, C:0.15-0.39, P<0.0001) (Table 4).

**Table 4.**
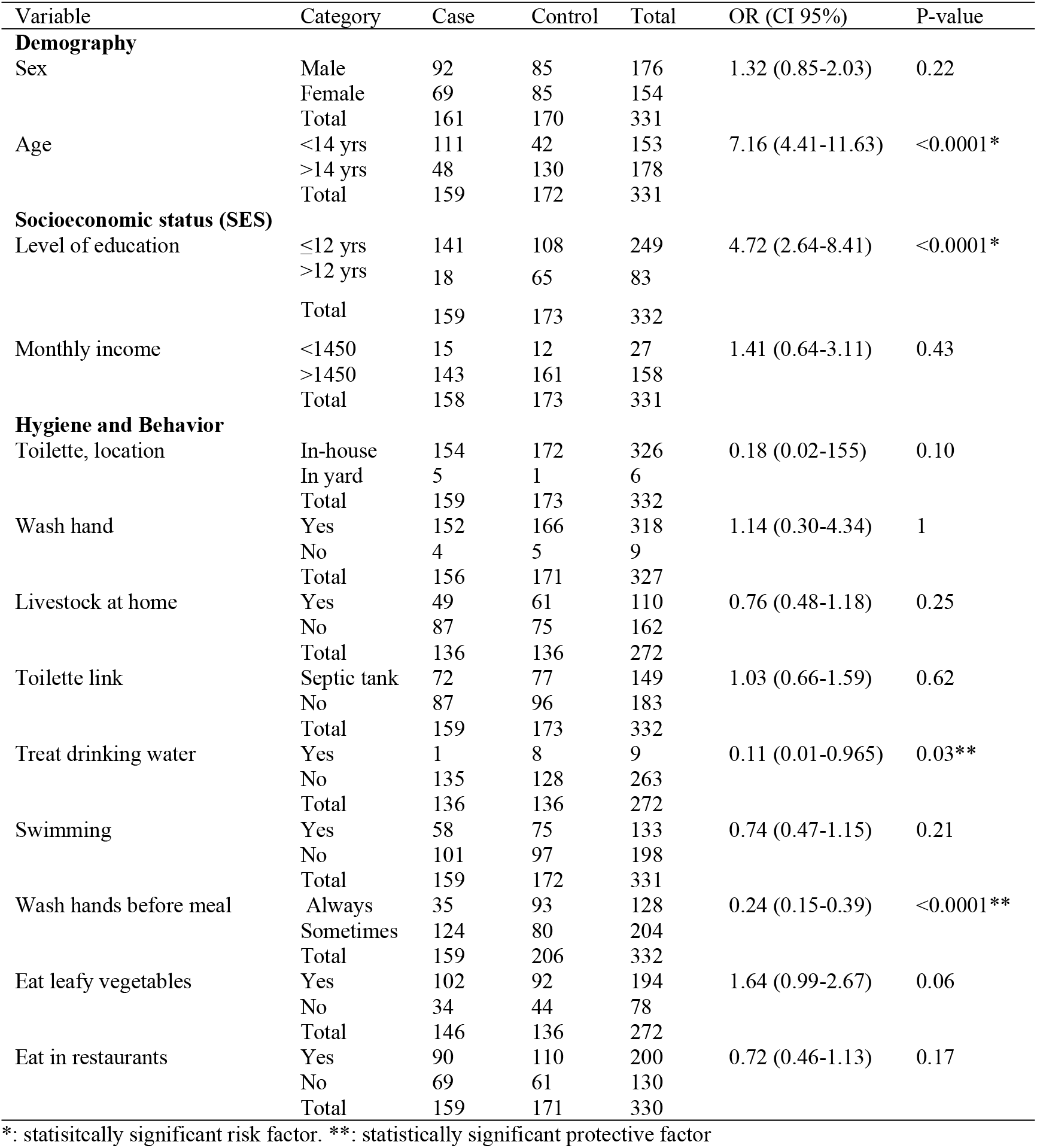
Demographic data, clinical history and the risk factors associated with acquiring HAV infection in the study group; demography, SES, and hygiene and behavior of the HAV cases and controls.

## Discussion

To the best of our knowledge, the present study is the first to use genetic diversity indices and haplotype analysis networking to analyze HAV variation from clinical sample. All of the 34 sequenced samples in the present study proved to be of the sub-genotype IB, which is the predominantly circulating genotype in the Mediterranean region from both clinical and environmental samples such as Spain, Jordan, Egypt and Turkey [25–28].

The maximum likelihood phylogenetic tree of the 23 HAV-IB sequences in the present study with other 28 HAV sequences retrieved from the gene bank, identified at least four clusters, but with weak bootstrap values. Although most (78%) of the samples isolated in 2015 and 2016 distinctively clustered in clade III, four Palestinian HAV-IB samples isolated between the years 2014 and 2016 intermixed with other HAV-IB isolates from Turkey, Spain, South African, Iran and Uruguay in clusters I. One isolate clustered uniquely by its own (Figure 1). Similar results had been described in a Bulgarian study which reported the splitting of HAV-IB into several clusters with few cases intermixing between Bulgarian and European isolates while others formed a unique Bulgarian cluster [4]. Furthermore, Wang *et al.*, (2013) delineated HAV isolates from the same area and during the same period to have clustered to several closely related lineages with low genetic diversity, suggesting either they possess a fitness advantage in the region, or an endemic transmission of closely related strains circulating in the neighboring regions [29]. The four Palestinian isolates in cluster I-and the lonely clustering isolate may have originated from travelers or food imported from HAV endemic areas, whereas, cluster III that consisted of purely Palestinian isolates indicates endemic HAV genotypes in Palestine. In addition to the weak bootstrap values, the genetic diversity indices such as total nucleotide diversity per site (π), average number of nucleotide differences between any two sequences (k), and neutrality indices supported the low population nucleotide diversity among HAV sequences.

The negative values of Tajima’s D and Fu-Li’s F indicate the amount of nucleotide variation observed (π) between HAV isolates is much less than expected (θ) which means low nucleotide variation. HAV clusters I and III with relatively high negative values for Tajima’s D and Fu’Li’s F statistics and low Hd (0.69) may have been subjected to recent population expansion or selection variation events that reduce genetic diversity such as selective sweep or bottleneck. This low variation between HAV clusters was further supported by the equal values of nucleotide diversity (π) in almost all cluster and the same applied to the average number of nucleotide differences (k). On the other hand, signs of polymorphism and nucleotide variation can be discerned in cluster IV which had positive Tajima’s D value (0.17). This cluster consisting of European HAV isolates can be argued to have begun a balancing selection where cluster IV is on the brink of developing into a distinct population (Table 1). Such data had been supported by recent published data from Bulgarian study which showed that the sequence analysis of HAV-IA subgenotypes were either identical or showed very few (1 to 4) nucleotide variations [30].

The fixation indices such as Fst confirmed genetic differentiation between the four HAV probable population with minimal migration among population reducing gene flow (Nm) between the cluster (Fst > 0.25 and Nm<1).

The total number of haplotypes produced by the haplotype analysis was 28. Indeed, several haplotypes (4-9) were observed in each phylogroup. This proves that despite low genetic diversity indices, haplotypes networking can be a good surrogate for unraveling the diversity among monomorphic population and even within the same phylogenetic group which is due to the fact that haplotype network analysis is based on detecting the single nucleotide variation (SNV) among studied sequences making it a powerful tool to detect genetic and haplotype variation. Interestingly, our study showed that the number of haplotypes in the clusters containing the Palestinian isolates, I and III, were the highest with h= 9 and 8, respectively (Table 2). The high number of haplotypes in cluster III indicate that the Palestinian HAV isolates, despite being from a geographically confined area, are more diverse and heterogeneous than those from Europe (cluster IV). However, despite the high number of haplotypes among our isolates, in comparison to the European cluster, the nucleotides variation is very low (1-2nts) which might be the result of a common origin coming from recent population expansion or selective sweep or genetic bottleneck.

Despite the high number of Palestinian haplotypes (red circles), but they are characterized by closely related viral haplotypes with 1-2 SNVs away from each other reflecting how close these haplotypes are. Cluster I is the most heterogeneous as it had the highest number of haplotypes (h=9), the haplotypes ≥2 SVNs apart from each other, and representing isolates from Asia, Europe, South America and Africa.

Moreover, the identical median-joining haplotype networks constructed by PopArt 1.7 and Network 10.0 (Figure 3) supported the low-bootstrap value phylogroups created by the maximum likelihood phylogenetic tree using MEGA version X (Figure 2) and running parallel with genetic and haplotype diversity parameters calculated by DnaSP ver. 6.12.03.

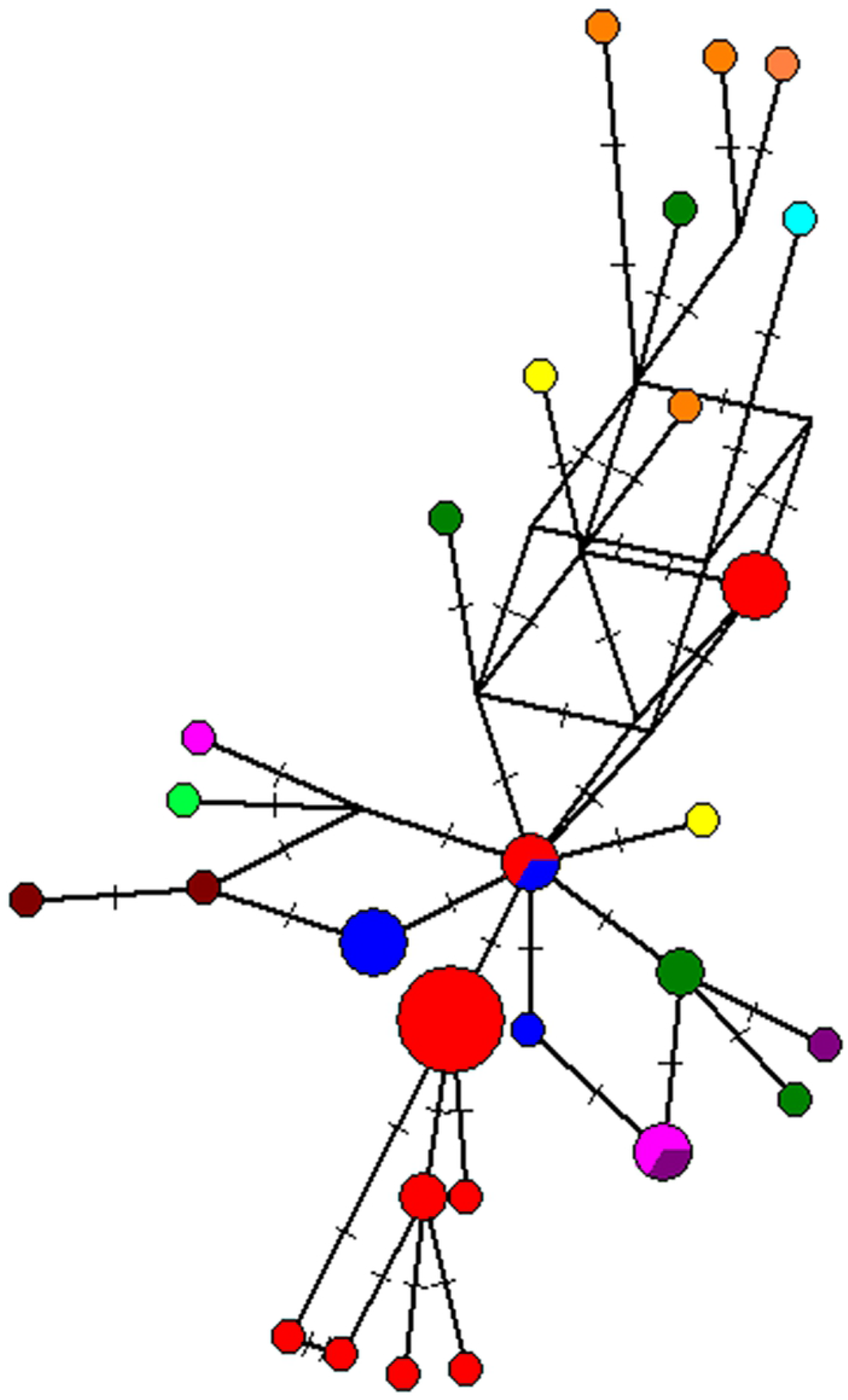

It is noteworthy, that even though a variety of genetic and haplotype diversity indices and several genetic differentiation parameters were calculated and networks and trees constructed, some variation may have been missed due to different reasons such as missing nucleotide during sequencing and insufficient number of sequences from study area and from Genbank.

In this study, young age, low level of parent’s education significantly increased the odds of HAV infection (OR>1, P<0.05), while the significantly low odds (OR<1, P<0.05) with the increase of hand washing before meal and treating of drinking water indicated the decrease of HAV infection suggesting these two variables are protective factors. On the contrary, Koroglu et al., (2017) reported that water and sanitation were not significant risk factors for HAV infection, whereas, gross domestic product (GDP), gross national income (GNI), and the human development index (HDI) were all highly associated with HAV infection rate in the Middle East and North Africa [6]. In addition, Hayajneh et al., (2015), reported that the incidence rate of HAV infection in Jordan decreased due to increase in level of maternal education, use of processed bottled drinking water and good sanitation practice [31–33]. Our results were in congruence with other studies that reported personal hygiene (hand washing before food preparation, cooking, eating and after defecation), living on crowded campuses, drinking unprocessed water, were the major risk factors of HAV infections [34–37]. In addition, political conflict in Palestine and the occupation of the West Bank and Gaza Strip, Palestine as well as the wars in neighboring Middle East countries usually, disturb the environmental and health condition and commonly increase the spreading of infectious disease outbreaks, including HAV, not only in source countries but also in neighboring regions that host refugees.

**In conclusion**, this study confirmed that based on VP3/VP1 junction region of the HAV genome; HAV possesses low genetic variation and nucleotide diversity, albeit haplotype network analysis revealed haplotype variation among the Palestinian sequences. Future studies based on multiple targets or full genome sequences are expected to give full picture on the genetic and haplotype variation. Our study reconfirmed that age and parent’s level of education as HAV risk factors, while hand washing and treating drinking water as protective factors.

